# Variation in rural African gut microbiota is strongly correlated with colonization by *Entamoeba* and subsistence

**DOI:** 10.1101/016949

**Authors:** Elise R Morton, Joshua Lynch, Alain Froment, Sophie Lafosse, Evelyne Heyer, Molly Przeworski, Ran Blekhman, Laure Segurel

**Affiliations:** Department of Genetics, Cell Biology, and Development, University of Minnesota, Minneapolis, MN, 55455; Eco-anthropology and ethnobiology, UMR 7206, CNRS-MNHN-University Paris 7 Diderot; Department of Biological Sciences, Columbia University, New York, NY 10027

## Abstract

The human gut microbiota is impacted by host nutrition and health status and therefore represents a potentially adaptive phenotype influenced by metabolic and immune constraints. Previous studies contrasting rural populations in developing countries to urban industrialized ones have shown that industrialization is strongly correlated with patterns in human gut microbiota; however, we know little about the relative contribution of factors such as climate, diet, medicine, hygiene practices, host genetics, and parasitism. Here, we focus on fine-scale comparisons of African rural populations in order to (i) contrast the gut microbiota of populations inhabiting similar environments but having different traditional subsistence modes and either shared or distinct genetic ancestry, and (ii) examine the relationship between gut parasites and bacterial communities. Characterizing the fecal microbiota of Pygmy hunter-gatherers as well as Bantu individuals from both farming and fishing populations in Southwest Cameroon, we found that the gut parasite *Entamoeba* is significantly correlated with microbiome composition and diversity. We show that across populations, colonization by this protozoa can be predicted with 79% accuracy based on the composition of an individual's gut microbiota, and that several of the taxa most important for distinguishing *Entamoeba* absence or presence are signature taxa for autoimmune disorders. We also found gut communities to vary significantly with subsistence mode, notably with some taxa previously shown to be enriched in other hunter-gatherers groups (in Tanzania and Peru) also discriminating hunter-gatherers from neighboring farming or fishing populations in Cameroon.

**Author Summary:** The community of microorganisms inhabiting the gastrointestinal tract plays a critical role in determining human health. It’s been hypothesized that the industrialized lifestyle, marked by a diet rich in processed foods, higher use of antibiotics, increased hygiene, and exposure to various chemicals, has altered microbiota in ways that are harmful. Studies have addressed this by comparing rural and industrialized populations, and have found that they systematically vary in their gut microbiome composition. Nevertheless, the relative influence of host genetics, diet, climate, medication, hygiene practices, and parasitism is still not clear. In addition, microbial variation between nearby human populations has not been explored in depth. Moreover, The World Health Organization estimates that 24% of the world’s population, concentrated in developing countries, is infected with gut parasites. Despite this, and evidence for direct interactions between the immune system and both gut parasites and bacteria, we know relatively little about the relationship between gut helminths, protozoa, and bacteria. In our study, we aimed to address some of this complexity. To do so, we characterized the gut microbial communities and parasites from Pygmy hunter-gatherer and Bantu farming and fishing populations from seven locations in the rainforest of Southwest Cameroon. We found that both subsistence mode and the presence of the gut protozoa, *Entamoeba*, were significantly correlated with microbiome composition. These findings support previous studies demonstrating diet is an important determinant of gut microbiota, and further show that this pattern holds true at a local scale, in traditional societies inhabiting a similar environment. Additionally, we show a significant relationship between a common human parasite (*Entamoeba*) and gut bacterial community composition, suggesting potential important interactions between the immune system, gut bacteria, and gut parasites, highlighting the need for more hierarchical cross population studies that include parasitism as potential factor influencing gut microbiota dynamics.

## Introduction

Humans and gut microbiota, the community of microorganisms inhabiting the gastrointestinal tract, have evolved in close association with each other for millions of years. As a result, humans depend on these microbes for acquisition of key nutrients from food, shaping the immune system, and providing protection from opportunistic pathogens (1-3). Despite considerable plasticity in the structure and composition of an individual’s gut microbiota (4), significant correlations between characteristics of the microbiome and host genotype (5-8), exposure to maternal microbiota (9), and patterns of disease (10, 11) suggest that the human microbiome represents a potentially adaptive phenotype with important implications for human health.

Since the Neolithic revolution about 12,000 years ago, human populations have started to diversify their dietary regimes, resulting in the contrasted subsistence modes known today. This major cultural transition has created metabolic constraints as well as novel pressures by pathogens due to the proximity of livestock and the increased density of populations. Such cultural and environmental differences among populations have resulted in physiological adaptations that can be detected in our genome (12-14) and have likely affected the community dynamics of our gut microbial ecosystem. Dietary changes have been shown to facilitate rapid changes in gut microbiota (4); however, the roles of habituation (over a lifetime) versus host adaptation (across generations) in these broader patterns are unclear. Understanding the longterm interaction that took place between the dietary specialization of populations and their gut microbiomes is therefore of great interest, notably to understand and predict the effect of recent and rapid changes in lifestyle and food on human health.

Nevertheless, to date, microbiome studies have mostly focused on industrialized populations. Of the few studies that have included a more diverse array of populations, most have contrasted urban populations in highly industrialized countries to populations in developing countries (15-19), or populations having both contrasted diet and occupying distinct climates (20, 21). Such designs do not allow the respective influences of the many factors coupled to geography such as diet, climate, hygiene, parasitism, and host genetics on microbiome variation to be disentangled. While some specific changes in microbial communities have been linked to components of human dietary regimes (4, 19, 22), urbanization levels (23, 24), hygiene practices (15), and the use of antibiotics (23, 25), the effect of other environmental or host-related factors is not clear. Notably, we do not know the extent to which the observed loss of microbial diversity of the human gut microbiome in urban industrialized populations (15-18, 20, 23) is attributable to their dietary specialization, differences in pathogen/parasite exposure, or other environmental factors (26). This loss of microbial biodiversity is a public health concern, as it may reflect a perturbed ecosystem associated with multiple diseases (27, 28).

In addition to the loss of microbial diversity, developed countries nearly ubiquitously present a marked decrease in the prevalence of human gut parasites (29). Although it is estimated that 3.5 billion people worldwide are infected with some parasite (protozoan or helminth) (30), studies assessing their role in shaping gut microbiota are limited (31). Yet throughout evolution, gut microbes and gut-dwelling parasites have co-inhabited the human gastrointestinal tract (32), and community dynamics are likely determined by current and past interactions (both during an individual’s lifespan and throughout evolutionary history) between microbiota, protozoa, helminths, and the host immune response (33). For example, it has been shown that direct competition by commensal microbes can provide protection from invading pathogens, and a disturbance to the natural microbiota can effectively result in increased susceptibility to pathogens and/or parasites (34, 35). There is also substantial evidence that these interactions are essential for the development of a healthy immune system, and that the underlying cause of the increased incidence of autoimmune disorders in industrialized countries is the absence of exposure to pathogens and parasites early in life (the “hygiene hypothesis”) (36, 37). In this context, it is important to evaluate the potential role of protozoa and helminths in shaping gut microbiota composition and structure.

Here, we focus on fine-scale comparisons of African rural populations with contrasting modes of subsistence but similar local environment and urbanization levels, and either shared or distinct genetic ancestry. Our objective is to better understand the relative influence of diet, host genetics, and parasitism on human gut microbiota composition and structure. We focus on populations from Cameroon for which a diversity of subsistence modes coexist in a restricted geographical area and a shared ecosystem (i.e., the tropical rainforest). We include individuals from hunter-gatherer populations (which are referred to in this manuscript as Pygmy to distinguish their genetic ancestry), Bantu farming populations and Bantu fishing populations, all living in a rural environment. These populations are almost entirely self-sufficient in food; their primary source of energy comes from cassava (*Manihot esculenta*), and fish or meat provides the main source of protein. Animal food production for these populations has been estimated to be high compared to elsewhere in Cameroon or Africa (38). To account for recent changes in diet, we evaluated current dietary regimes using dietary surveys. We also assessed parasitism status by direct observations of fecal samples under the microscope. The focus on populations living in the tropical rainforest is complementary to previous African populations sampled: the Hadza hunter-gatherers and a population from Burkina Faso, living in the East and West African tropical savanna, respectively (15, 18) and a population from Malawi living in a relatively dry subtropical area of East Africa (16). Here, in addition to comparing the gut microbiota of human populations with limited geographic separation and contrasting subsistence modes, we aimed to characterize the relationship between gut microbial communities and various intestinal parasites.

## Results

### Description of the samples: host population, diet, and parasites

We analyzed 64 individuals in seven different villages in Southwest Cameroon, corresponding to 20 hunter-gatherers, 24 farmers, and 20 individuals from a fishing population (see Figure 1 and Table S1). The average age of study participants ranges between 26 and 78 years, with an average age of 50 years. The Pygmy hunter-gatherers diverged from the other Bantu populations about 60,000 years ago (39, 40) and the farming subsistence mode likely started over the last 5,000 years (41). The sampled populations therefore not only have contrasted subsistence modes, but also have different genetic backgrounds.

**Figure 1.**
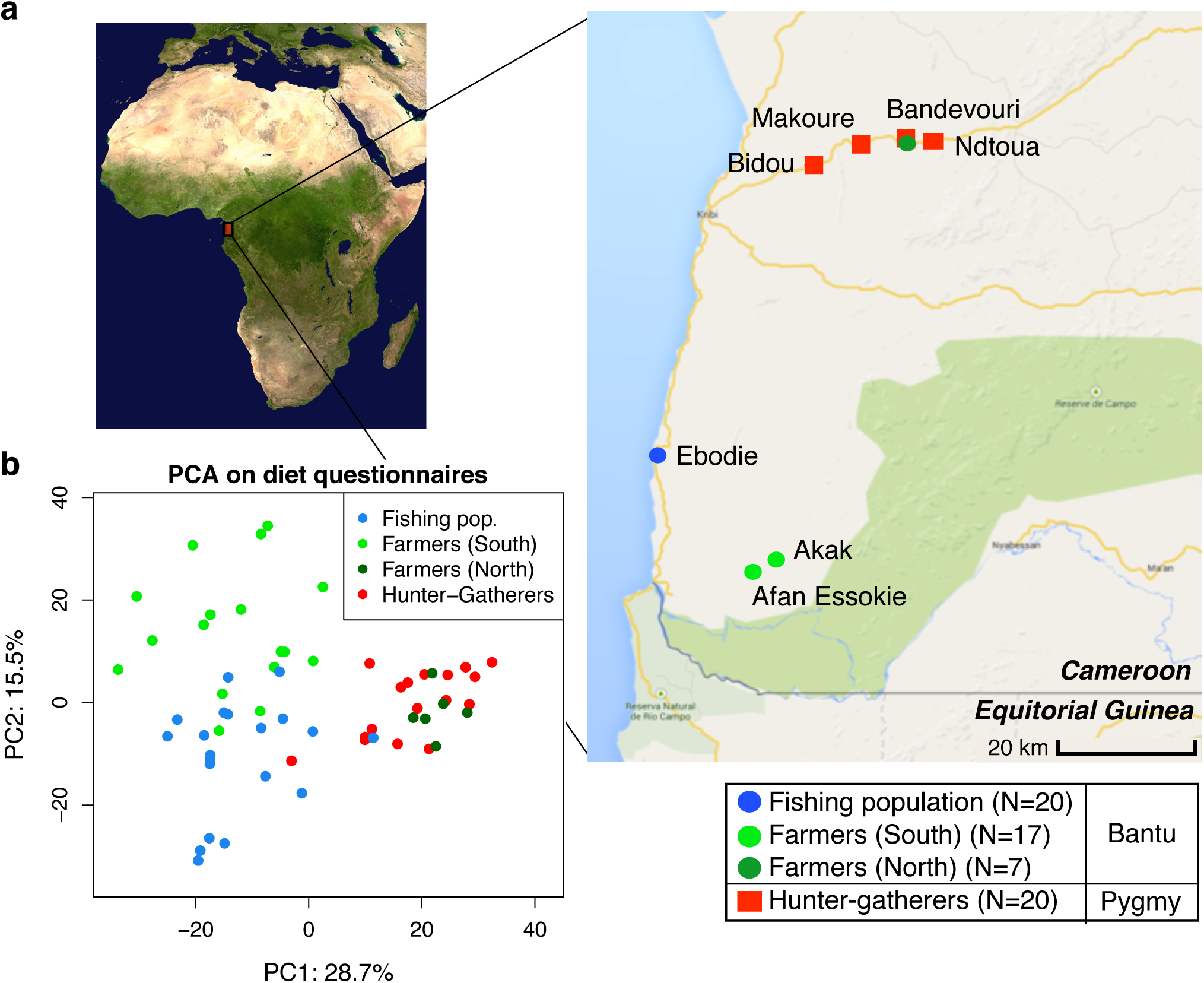
(a) Map showing the geographic locations of the villages sampled in Southwest Cameroon, the number of samples (N) collected for each subsistence group (the fishing population, farmers from the South, farmers from the North, and hunter-gatherers), and their genetic ancestry (Bantu or Pygmy). (b) Principle Components Analysis based on dietary questionnaires for all 64 individuals. The first two principal components (PC1 and PC2) are shown, with the amount of variation explained reported for each axis.

We chose these populations because previous work done in 1984-1985, based on nutritional questionnaires and isotopes analyses, showed they had distinct diets (38, 42). We performed new nutritional frequency surveys to assess how diet had changed during the past 30 years (see Table S1). Interestingly, the amount of meat in the hunter-gatherers’ diet has substantially decreased, reflecting the lower abundance of wild game in the forest reserve and the hunting ban applied for some species. In contrast, the consumption of fish has increased in inland populations (especially in farmers), due to the construction of new roads connecting the coastal and inland populations. Similar to the results from 1984-1985, the farmers eat less starchy foods (cassava) than hunter-gatherers and individuals from the fishing population (p = 0.005 and 0.017, respectively; Mann-Whitney U test). A principal component analysis on all dietary components revealed roughly three clusters corresponding to the three dietary regimes, with the first axis distinguishing hunter-gatherers from the others, and the second axis separating the farming and fishing populations (see Figure 1b). The one exception to this pattern concerns the farmers from the North (living along the same road as the hunter-gatherers), who cluster with the hunter-gatherers. Based on a permutation test (10,000 permutations), the Euclidean distance between the hunter-gatherers (grouped with the North farmers) and the South farmers and fishers, respectively, is significant (p < 0.0001 in both cases), whereas between South farmers and fishers it is not (p=0.3). Therefore, in our analyses of subsistence we consider the North and South farmer populations separately.

In addition to dietary questionnaires, we assessed the nutritional status of individuals by measuring their BMI (Body Mass Index) (see Table S1). Twenty percent of the Pygmy hunter-gatherers were underweight (BMI < 18) whereas 12%, 0%, and 4% of the South farmers, North farmers, and individuals from the fishing population were, respectively. Conversely, 0% of hunter-gatherers were overweight (BMI > 25) while 12%, 14% and 26% of individuals in the other groups were, respectively. This likely reflects the difference in socio-economical status and access to medicine between these populations. Subsistence (as defined by the four following groups: hunter-gatherers, farmers from the North, from the South and fishers) was significantly correlated with BMI in a linear regression model (p = 0.026), but not using a Pearson Chi-square test (p = 0.25).

We assessed the intestinal parasitism of individuals by direct observation of their fecal samples under the microscope and detected the presence of *Entamoeba* cysts, as well as eggs of *Ascaris*, *Trichuris*, and *Ancylostoma* (see Table S1 and Figure S1). Overall, 89% of hunter-gatherers, 76% of farmers from the South, 100% of farmers from the North, and 58% of individuals from the fishing population were infected by at least one of these organisms. Regarding *Entamoeba*, 37%, 41%, 57% and 16% of individuals were infected in each population, respectively. Although the presence of *Entamoeba* was not significantly correlated with subsistence (p = 0.18; linear regression model), the reduced rate of parasitism in the fishing population most likely reflects their higher level of hygiene and increased access to medicine. However, further studies are needed to examine the effects of medication on parasitism in these populations. Based on Pearson’s Chi-squared tests, we found that there were no statistically significant relationships between *Entamoeba* and any of the covariates tested (including sex, age, BMI, subsistence, genetic ancestry, location, or dietary components; p > 0.1). However, a linear regression analysis found a correlation between *Entamoeba* and age (p = 0.019), but not with the other factors (Figure S10).

### Characterization of microbiome composition

The fecal microbiota of 69 samples (including 5 biological replicates) were characterized by sequencing of the V5-V6 region of the bacterial 16S ribosomal RNA with the Illumina MiSeq technology. The dataset was rarefied to 50,000 reads/sample (see Figure S2), and reads were clustered into 5039 operational taxonomic units (OTU) at 97% identity.

The five biological replicates (sampling of the same individual few days apart, see Table S1) allowed us to compare the microbial differences within individuals to those between individuals. We assessed differences in gut communities by calculating UniFrac distances, a phylogenetic based distance metric, which when weighted, accounts for relative abundance of taxa (43). Because both weighted and unweighted metrics capture different aspects of microbial diversity (43), we included both types of analyses in the manuscript. We found that the average UniFrac distance between replicates of the same individual was lower than between individuals (although only statistically significant for the unweighted distances: p = 0.003; one-sided Mann-Whitney U test; see Figure S3).

We used PERMANOVA analysis to separately test for associations between microbiome composition (OTU abundances) and age, sex, BMI, parasitism, location, subsistence, and ancestry, the latter three being nested (see Table S2). We found that the presence of *Entamoeba*, location, subsistence, and ancestry were each significantly associated with variation in microbiome composition (p = 0.0001, 0.01, 0.003 and 0.01, respectively; Table S2), whereas the other factors were not. To further characterize patterns of variation that account for phylogenetic relationships of community taxa, we also performed a PERMANOVA analysis on both weighted and unweighted UniFrac distance matrices. Congruent with our previous results, we found that the presence of *Entamoeba* was the most significant variable for both weighted and unweighted UniFrac distances (p = 0.007 and 0.0001, respectively; Table S2). *Entamoeba* infection also provided the strongest separation along the primary axis of variation of the multidimensional scaling plots (Figure 2a and Figure S4a). Subsistence and location were both determined to be significant based on unweighted UniFrac distances (p = 0.0003 and p = 0.002, respectively), but not weighted (p = 0.14 and p = 0.29, respectively). Because unweighted UniFrac distances assign increased weight to rare taxa, this suggests that less abundant taxa are more important in describing differences between the microbiomes across subsistence modes and locations. Furthermore, subsistence provided only weak visual separation along the first two axes of variation for both metrics (Figure S4b-c).

**Figure 2.**
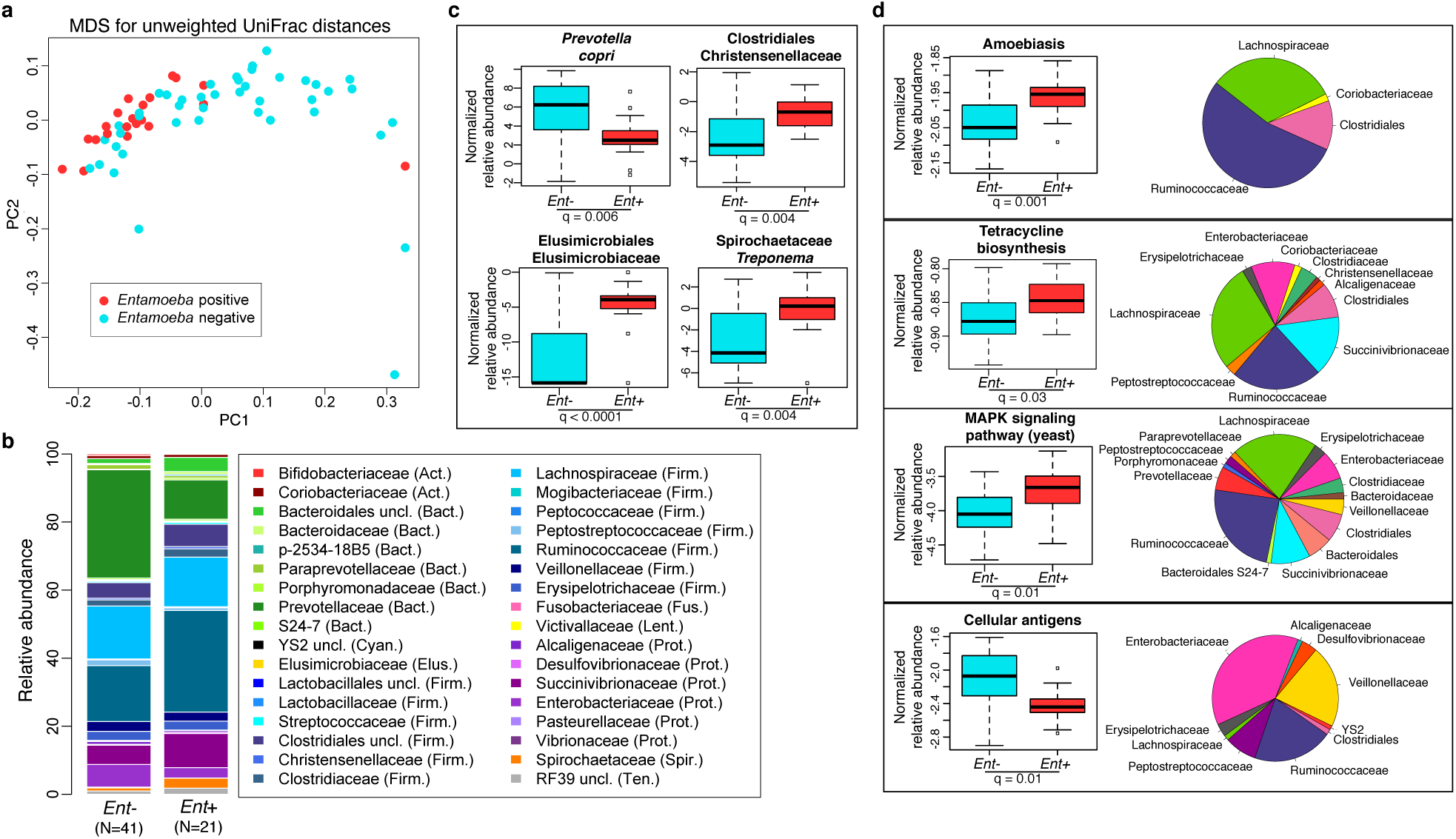
Relationship between the presence of *Entamoeba* (*Ent*- or *Ent*+) and fecal microbiome composition. (a) Multidimensional Scaling plot of unweighted UniFrac distances colored by *Entamoeba* presence or absence. The first two principal components (PC1 and PC2 are shown). (b) Summary of the relative abundance of taxa (>= 0.1% in at least 4 individuals) for *Ent*- and *Ent*+ individuals color coded by phylum (Actinobacteria (Act.) = red, Bacteroidetes (Bact.) = green, Cyanobacteria (Cyan.) = black, Elusimicrobia (Elus.) = gold, Firmicutes (Firm.) = blue, Fusobacteria (Fus.) = pink, Lentisphaerae (Lent.) = yellow, Proteobacteria (Prot.) = purple, Spirochaetes (Spir.) = orange, and Tenericutes (Ten.) = gray). The number of individuals (N) in each population is indicated below the bars. (c) Normalized relative abundance of four taxa significantly associated with *Entamoeba* presence/absence in an ANOVA as well as in the random forest classifier model (q < 0.05). (d) Normalized relative abundance of four KEGG metabolic pathways significantly associated with *Entamoeba* status in an ANOVA (q < 0.05 using the most abundant (>= 0.4% in at least one group) KEGG (Level 3) pathways) (left panel) and the relative contributions of each taxon for each pathway (right panel).

### Relationship between parasitism and the microbiome

Because of the significant relationship between the presence of *Entamoeba* with patterns in variation in the gut microbial communities found in all populations, we further investigated the relationship between its presence and microbiota composition (Figure 2). As it is difficult to distinguish between the opportunistic pathogenic species (E. *histolytica*) and the strict commensal (E. *dispar*) by microscopy alone, we were unable to characterize this organism at the species level. However, only two of the sampled individuals self-reported to be suffering from diarrhea (one positive with *Entamoeba*, the other negative), suggesting that individuals with *Entamoeba* were not experiencing symptomatic amoebiasis. Previous studies showed that when both species are common in a population, there is a higher prevalence of *E. dispar* than *E. histolytica* and the majority of infections by *E. histolytica* are asymptomatic (44).

We first verified that the presence of *Entamoeba* was significantly associated with the gut microbiome including age, sex, BMI, and subsistence, ancestry, or location as covariates (p=0.0005, p=0.0003, and p=0.0001, respectively; PERMANOVA for unmerged OTUs). At the phylum level, we found that 7 of the 13 phyla represented are significantly different between individuals that harbored *Entamoeba* and those that did not (*Ent*+ and *Ent*-, respectively), with most phyla (except Bacteroidetes) occurring at a higher relative abundance in *Ent*+ individuals (see Table 1). When looking at individual taxa, based on an ANOVA, we also identified a number of notable differences between *Ent*+ and *Ent*- individuals (Figure 2b-c, Tables S3 and S4), and we found that eighteen of the 93 most abundant taxa (present at ≥ 0.1% in at least 4 individuals) differed significantly in their relative abundance between *Ent*+ and *Ent*- individuals (FDR q < 0.05, after Benjamini-Hochberg correction for the number of taxa analyzed (45)). To ensure that these relationships were not due to other factors, we included age, sex, BMI, and either subsistence, location, or ancestry as covariates in the model and found that although the q-values changed slightly, all were still significant (q < 0.05).

**Table 1.** Differences in abundances of gut microbiota across *Entamoeba* and subsistence. (a) Frequency (in %) of phyla for *Entamoeba* negative (*Ent*-) and positive (*Ent*+) individuals and for the four subsistence groups (Fis = Fishing population; Far(S) = Farmers from the South; Far(N) = Farmers from the North; HG = Hunter-gatherers). (b) Frequency (in %) of specific taxa of interest previously associated with geography in the four subsistence groups. FDR Q-values are based on an ANOVA (controlling for subsistence and *Entamoeba*, in (a) and (b) respectively), after Benjamini-Hochberg correction for the number of tests. The first letters in parenthesis indicate to which phylum each taxa belongs (Act. = Actinobacteria, Bact. = Bacteroidetes, Firm. = Firmicutes, Prot. = Proteobacteria, and Spir. = Spirochaetes).

| Phylum (>=0,1% in at least 4 ind) | Effect of *Entamoeba* |  |  | Effect of subsistence |  |  |  |  |
| --- | --- | --- | --- | --- | --- | --- | --- | --- |
|  | Ent - | Ent + | q-val | Fis | Far(S) | Far(N) | HG | q-val |
| **Actinobacteria** | 1.4 | 1.0 | 0.866 | 2.1 | 0.9 | 0.9 | 0.8 | 0.148 |
| **Bacteroidetes** | 35.3 | 18.6 | 0.008 | 33.4 | 31.7 | 26.7 | 26.0 | 0.664 |
| **Cyanobacteria** | 0.17 | 0.24 | 0.026 | 0.10 | 0.26 | 0.18 | 0.24 | 0.148 |
| **Elusimicrobia** | 0.03 | 0.10 | 2E-05 | 0.01 | 0.08 | 0.23 | 0.05 | 0.148 |
| **Euryarchaeota** | 0.03 | 0.09 | 2E-04 | 0.02 | 0.04 | 0.08 | 0.07 | 0.165 |
| **Firmicutes** | 47.0 | 60.8 | 0.026 | 49.5 | 55.3 | 58.0 | 47.8 | 0.576 |
| **Fusobacteria** | 0.15 | 0.02 | 0.401 | 0.13 | 0.21 | 0.01 | 0.01 | 0.351 |
| **Lentisphaerae** | 0.19 | 0.15 | 0.074 | 0.09 | 0.26 | 0.29 | 0.14 | 0.220 |
| **Proteobacteria** | 13.8 | 14.1 | 0.701 | 12.4 | 7.2 | 8.3 | 23.0 | 0.067 |
| **Spirochaetes** | 0.9 | 2.9 | 0.001 | 1.1 | 2.6 | 3.2 | 0.7 | 0.494 |
| **Tenericutes** | 1.0 | 1.9 | 0.008 | 1.1 | 1.5 | 1.9 | 1.0 | 0.220 |
| **Verrucomicrobia** | 0.04 | 0.16 | 0.272 | 0.02 | 0.04 | 0.23 | 0.12 | 0.165 |

Table 1. (a) Frequency (in %) of phyla for *Entamoeba* negative (*Ent*-) and positive (*Ent*+) individuals and for the four subsistence groups (Fis = Fishing population; Far(S) = Farmers from the South; Far(N) = Farmers from the North; HG = Hunter-gatherers).
| Specific taxa of interest | Fis | Far(S) | Far(N) | HG | q-val |
| --- | --- | --- | --- | --- | --- |
| (Act.) *Bifidobacterium all species* | 0.86 | 0.15 | 0.11 | 0.06 | 6E-05 |
| (Act.) *Bifidobacterium adolescentis* | 0.51 | 0.07 | 0.00 | 0.01 | 0.003 |
| (Bact.) (*Prevotellaceae*) *Prevotella all species* | 30.8 | 26.2 | 19.2 | 20.2 | 0.590 |
| (Bact.) *Bacteroides all species* | 0.22 | 0.38 | 0.25 | 0.84 | 0.692 |
| (Bact.) *Bacteroidales unclassified* | 0.67 | 3.14 | 4.56 | 2.44 | 5E-04 |
| (Firm.) *Lachnospiraceae unclassified* | 7.7 | 9.8 | 11.3 | 5.7 | 0.027 |
| (Firm.) *Ruminococaceae Ruminococcus all species* | 0.81 | 1.20 | 1.14 | 1.29 | 0.178 |
| (Firm.) *Veillonellaceae unclassified* | 0.22 | 0.35 | 0.67 | 0.21 | 0.021 |
| (Prot.) *Succinivibrio all species* | 5.7 | 3.3 | 2.8 | 9.7 | 0.025 |
| (Prot.) *Ruminobacter all species* | 0.07 | 0.10 | 0.02 | 3.74 | 0.021 |
| (Prot.) *Klebsiella all species* | 0.55 | 0.36 | 0.47 | 0.24 | 0.806 |
| (Prot.) *Salmonella all species* | 0.04 | 0.01 | 0.02 | 0.09 | 0.652 |
| (Spir.) *Treponema all species* | 1.1 | 2.6 | 3.1 | 0.7 | 0.806 |

These taxonomic signatures for *Entamoeba* status are so strong that its presence can be predicted with 79% accuracy using a Random Forest Classifier (RFC) model based on gut microbiome composition (p < 0.001; See Methods section Figure S5). Of the ten taxa identified as being the most important in their predictive power, all but *Prevotella stercorea* were significant in our ANOVA model (of which all are in higher abundance in *Ent*+ individuals except *Prevotella copri*). The reason for the association between *Entamoeba* and these microbes have yet to be identified, but it is noteworthy that the two most important taxa identified in the RFC model, Elusimicrobiaceae unclassified (uncl) and Ruminococcaceae uncl, include established endosymbionts of protists and common inhabitants of the termite gut (46). Furthermore, Ruminococcaceae uncl was shown to be enriched in Hadza as compared to Italians (18). Spirochaetaceae *Treponema*, the third most important taxon, include species that have been reported to inhabit the cow rumen, the pig gastrointestinal tract, and the guts of termites (47) and have been proposed as symbionts in the human “ancestral microbiome” (18, 20, 48). *Christensenellaceae*, the fourth most important taxon, was recently identified as being the most heritable taxon in an analysis of twins from the UK, and was shown to impact host metabolism (5). Two taxa in the order Bacteroidales, *Prevotella stercorea* and *Prevotella copri*, the seventh and eighth most important taxa, are the only ones occurring at significantly reduced abundance in infected individuals; *Prevotella* is an important genus of gut bacteria and is systematically underrepresented in Western microbiomes (15-18, 20, 26). While members of the Clostridia and Gammaproteobacteria are more abundant in infected individuals, the pattern for Bacteroidales is the opposite (see Figure 2b). *Oscillospira uncl* and Parabacteroides uncl, the ninth and tenth most important taxa, are associated with the rumen and human intestine, respectively.

Furthermore, when looking at the microbial diversity of *Ent*+ versus *Ent*- individuals, we found that the presence of *Entamoeba* is associated with a significant increase in alpha (intra-host) diversity using the Phylogenetic Distance Whole Tree metric (p = 1.03E-06; Welch’s t-test; Figure 3a), as well as using the Shannon and Simpson indices (p = 0.001 and p = 0.025, respectively; Welch’s t-test; Figure S6). Interestingly, although the alpha (intra-host) diversity of *Ent*+ individuals is significantly higher than *Ent*- individuals, the beta (inter-host) diversity (as estimated by both UniFrac distance metrics) reveals that gut communities across *Ent*+ individuals are more similar than across *Ent*- individuals (weighted and unweighted, p = 2.23E-06 and p < 2.2E-16; Welch’s t-test; Figure 3b and Figure S7). This could suggest that, as alpha diversity increases, there are fewer potential stable states for individual gut communities, or that the presence of *Entamoeba* drives changes in the microbiome (directly or indirectly through effects on the immune system) that are dominant over other factors.

**Figure 3.**
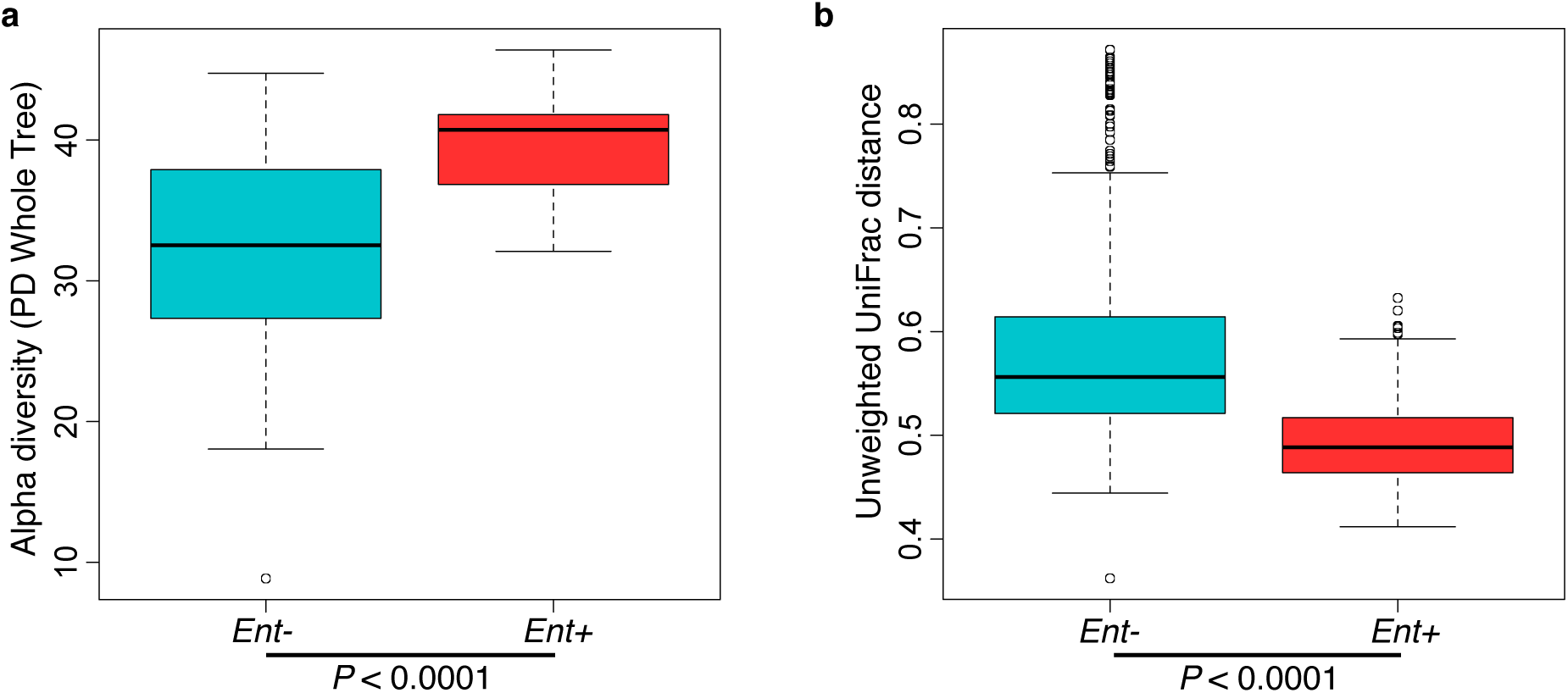
(a) Comparison of alpha diversity for *Entamoeba* negative (*Ent*-) and positive (*Ent*+) individuals using the phylogenetic distance whole tree metric. (b) Comparison of beta diversity within *Ent*-, within *Ent*+, and between *Ent*- and *Ent*+ individuals based on unweighted UniFrac distances. P-values are based on Welch’s t-test.

#### Relationship between specific taxa and microbial community diversity

Because of the striking relationship between the presence of *Entamoeba* and alpha diversity, we sought to identify any phyla for which abundance was significantly correlated with community diversity. To account for the effect of *Entamoeba*, we added it as a binary covariate to our ANOVA and identified 11 phyla that are significantly correlated with alpha diversity (FDR q < 0.05, after multiple test correction; see Figure S8). Although, as expected, the majority of these taxa increase in abundance with higher diversity, Bacteroidetes and Proteobacteria exhibit a decrease in relative abundance as alpha diversity increases. This negative relationship suggests that these taxa might be more competitive than others and drive down diversity.

#### Predicted metagenome

We used the KEGG (Kyoto Encyclopedia of Genes and Genomes) database (49) and the PICRUSt (Phylogenetic Investigation of Communities by Reconstruction of Unobserved States) pipeline (50) to predict abundances of pathways across individuals (see Figure S9). Many of these pathways are classified based on eukaryotic genes. However, homologues in prokaryotes could have related functions. Considering the 220 most abundant KEGG pathways (comprising ≥ 0.01% of all assigned reads in at least 4 individuals), we identified 19 pathways with significant differences in abundance between *Ent*+ and *Ent*- individuals (FDR q < 0.05 after Benjamini-Hochberg correction for the number of pathways tested; ANOVA; see Figure 2d and Table S6). Of these 19, of particular interest are an increase in amoebiasis (q = 0.001), biosynthesis of the antibiotic tetracycline (q = 0.03), and yeast MAPK signaling pathways (q = 0.01) in *Ent*+ individuals. These changes are largely attributed to Clostridiales and Ruminococcaceae, which occur at significantly greater abundance in *Ent*+ individuals (6.53% vs. 4.53%, q = 0.044; and 29.58% vs. 16.34%, q < 0.0001, respectively, Figure 2d). Interestingly, the Cellular Antigens pathway, potentially involved in host-microbe and microbe-microbe interactions, is more represented in the predicted metagenomes of *Ent*- individuals (q = 0.01; ANOVA). This pathway is predominantly attributed to members of the Enterobacteriaceae family, which was found to be twice as abundant in individuals lacking the parasite.

Finally, outside of an association between *Ancylostoma* and Bacteroidales uncl (q = 0.019; ANOVA), none of the other parasites tested (*Ascaris* and *Trichuris*) exhibited a significant association with any taxon, whether individually or as the number of all non-*Entamoeba* parasite types present. However, the overall composition seems to shift with the number of parasites (see Figure S11), and there is a significant increase in alpha diversity when three parasite species are present relative to just one (Figure S12).

### Relationship between subsistence mode and gut microbiota

#### Microbial community patterns across subsistence

Controlling for the effect of *Entamoeba*, subsistence mode was significantly correlated with patterns of gut microbiota (p = 0.004; PERMANOVA). To investigate the relationship between subsistence and microbiome community composition, we summarized microbial taxonomic composition across the four subsistence groups and their geographic locations (Figure 4a, Figure S13). At the phylum level, we found a moderately significant difference in the relative contribution of Proteobacteria across subsistence (q = 0.067, Table 1; ANOVA), with hunter-gatherers having a higher frequency than the fishing population, farmers from the South and the North (23% versus 12.4%, 7.2% and 8.3%, respectively), mirroring the higher frequency observed in the Hadza hunter-gatherers compared to Italians (18) and the higher frequency observed in traditional Peruvian groups (both hunter-gatherers and farmers) compared to US individuals. Based on an ANOVA, we also found that 8 of the most abundant taxa differed significantly across subsistence modes (see Figure 4b and Tables S3 and S5). Of particular interest is the genus *Bifidobacterium*, both *B. uncl* and *B. adolescentis*, which were found at higher abundance in the fishing population (means 0.30% and 0.51%, respectively) relative to all other populations (≤ to 0.11% and 0.07%, respectively; q = 0.0003 and q = 0.008; ANOVA). This genus is associated with a higher consumption of dairy products, a pattern also observed in a comparison of Italians to Hadza hunter-gatherers (18) and consistent with the occasional consumption of yogurt in the fishing population. We also found Bacteroidales uncl to occur at significantly lower relative abundance in the fishing population relative to the other three populations (0.7% vs. ≥ 2.4%; q = 0.003; ANOVA), an order of bacteria also identified as being less abundant in the Italians versus the Hadza (18). In contrast with other Firmicutes genera that tend to be in lower frequency in hunter-gatherers, we found the genus *Sarcina,* a synthesizer of microbial cellulose, to be only present in the hunter-gatherers (means of 0.69% compared to ≤ 0.07% in the other subsistence groups; q = 0.007; ANOVA). This genus was also found in higher frequency in traditional farming populations from Papua New Guinea as compared to US individuals (26). Finally, we found three members of the Lachnospiraceae family to be significantly different among populations, with *Ruminococcus uncl* and *Ruminococcus gnavus* being in lower frequency in hunter-gatherers (0.34% and 0.19%, respectively) compared to other populations (0.46-0.86% and 0.41-0.99%, respectively; q = 0.030 and 0.006; ANOVA). This family has been linked to obesity (51) in addition to protection from colon cancer attributable to their production of butyric acid (52). Importantly, although BMI did differ across subsistence modes (p = 0.026, reflecting a BMI significantly higher in fishers; linear regression model), we did not find a significant relationship between BMI and microbiota composition or diversity patterns (see Table S2).

**Figure 4.**
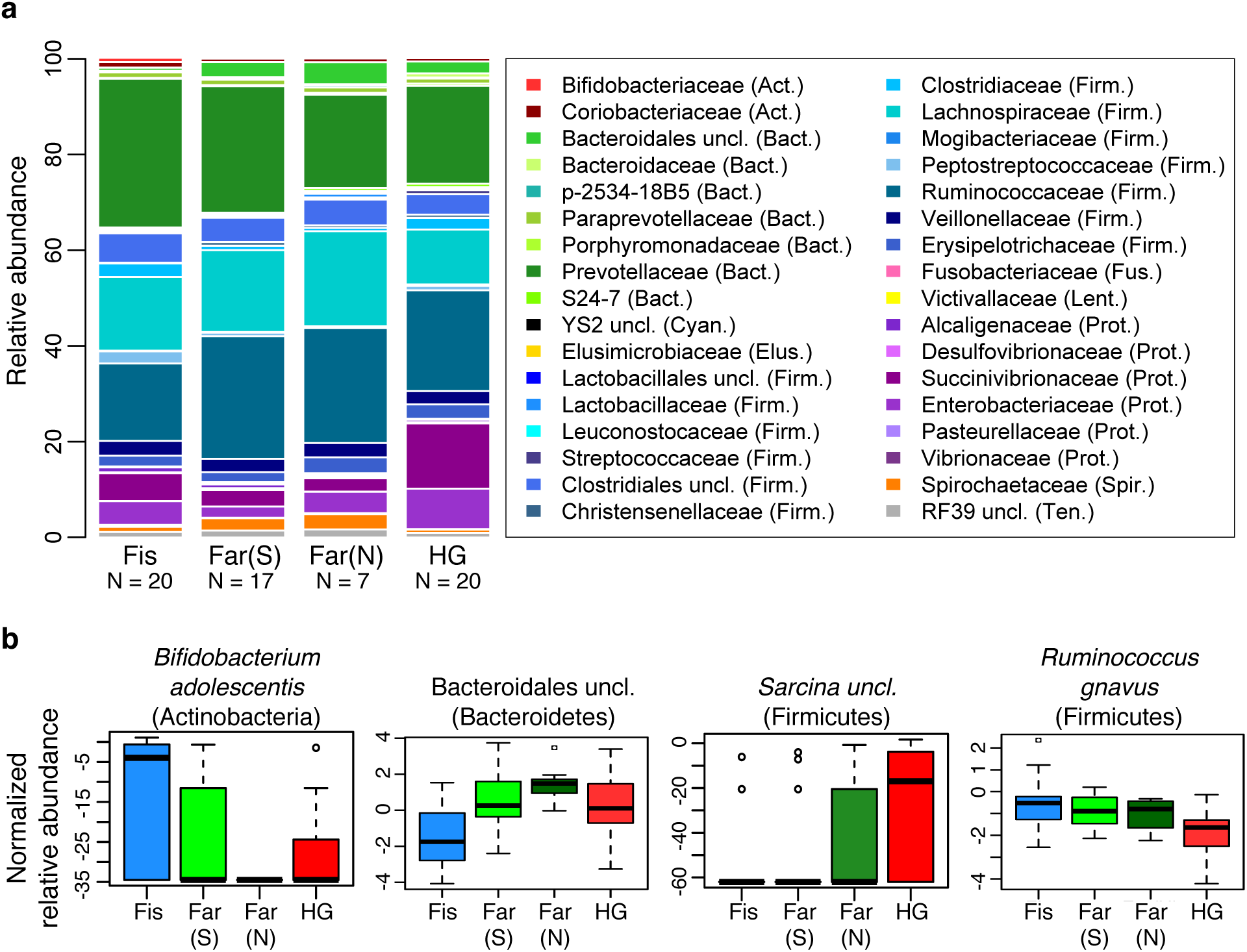
Relationship between subsistence modes and fecal microbiome composition. (a) Summary of the relative abundance of taxa (occurring at >= 0.1% in at least 4 individuals) for individuals across subsistence. Taxa are colored by phylum (Actinobacteria (Act.) = red, Bacteroidetes (Bact.) = green, Cyanobacteria (Cyan.) = black, Elusimicrobia (Elus.) = gold, Bacteroidetes (Bact.) = green, Cyanobacteria (Cyan.) = black, Elusimicrobia (Elus.) = gold, Firmicutes (Firm.) = blue, Fusobacteria (Fus.) = pink, Lentisphaerae (Lent.) = yellow, Proteobacteria (Prot.) = purple, Spirochaetes (Spir.) = orange, and Tenericutes (Ten.) = gray). The number of individuals (N) in each population is indicated below the bars. (b) Relative abundance of four taxa significantly associated with subsistence based on an ANOVA, q < 0.05. Fis = Fishing population; Far(S) = Farmers from the South; Far(N) = Farmers from the North; HG = Hunter-gatherers.

A random forest classifier (RFC) model for microbiome composition revealed an overall accuracy of 59% (p < 0.001) for predicting the four subsistence groups but varied widely across populations (see Table S7 and Figure S14a). The hunter-gatherer population was the most distinguishable such that the correct subsistence group was accurately predicted 85% of the time (versus 31% if predictions had been made by chance alone). Individuals of the fishing and South farming populations were predicted with 65% and 47% accuracy, respectively (versus 31% and 27% by chance), and the North farming population was never predicted correctly (versus 11% by chance, suggesting their microbiota are variable but share patterns with the other populations). Furthermore, incorrect assignments for individuals of the hunter-gatherer, farmers from the South and fishing populations were distributed evenly across all other subsistence groups, with the exception of farmers from the North, to which no individual was predicted to belong. In agreement with our ANOVA, the taxon identified as being the most important in distinguishing subsistence groups was *Bifidobacterium uncl* (see Figure 4b and Figure S14b), occurring at significantly higher frequency in the fishing population (q = 0.0003, Table S5). *Ruminococcus bromii*, important for degradation of resistant starch (53), was the second most important taxon, occurring at 0.01%, 0.01%, 0.15%, and 0.12% in the fishing population, farmers from the North, the South, and hunter-gatherers, respectively (q < 0.0001) (see Figure S14c). The third, fourth, fifth and eighth most important taxa include members of the Lachnospiraceae family, two of which were found to be significant based on an ANOVA (see above). When grouped together, taxa in this family are less abundant in the hunter-gatherers relative to other subsistence groups (11.3% vs. 15.6-19.6%, respectively), a difference significant only when comparing hunter-gatherers to both farmer populations. Finally, two species of the Succinivibrionaceae family, *Succinivibrio sp.* and *Ruminobacter sp.*, were also identified as being important taxa in the model, both of which were more abundant in the hunter-gatherers at 9.7% and 3.7%, respectively, vs. less than 5.7% and less than 0.1% for the other three subsistence modes (q = 0.068 and 0.057, respectively; see Figure S14c). These taxa, associated with the bovine rumen, were also found in higher frequency in the Hadza hunter-gatherers and traditional Peruvian populations (18, 20). Finally, only five of the top ten taxa identified in the random forest classifier model were determined to be significant in the ANOVA (see Figure S14b). This suggests that rather than an individual signature taxon, it is likely the pattern of abundances of multiple taxa that is important for predicting subsistence.

#### Diet and gut microbial diversity

We found the alpha (intra-host) diversity to be significantly lower in the fishing population than in farmers from the South and the North for the phylogenetic distance whole tree metric (p = 0.021 and p = 0.008, respectively; Welch’s t-test) and only compared to farmers from the North for the Shannon and Simpson metrics (p = 0.017, and 0.021, respectively; Welch’s t-test) (see Figure 5a and Figures S15-17). Interestingly, the pattern of beta diversity across subsistence modes using both unweighted and weighted UniFrac distance metrics also distinguishes the fishing population from both farmers, such that the within-group variation is significantly higher in the fishing and hunter-gatherer populations compared to both farmers (p < 0.001 for all relevant pairwise comparisons; Welch’s t-test) (see Figure 5b and Figure S17a-b).

**Figure 5.**
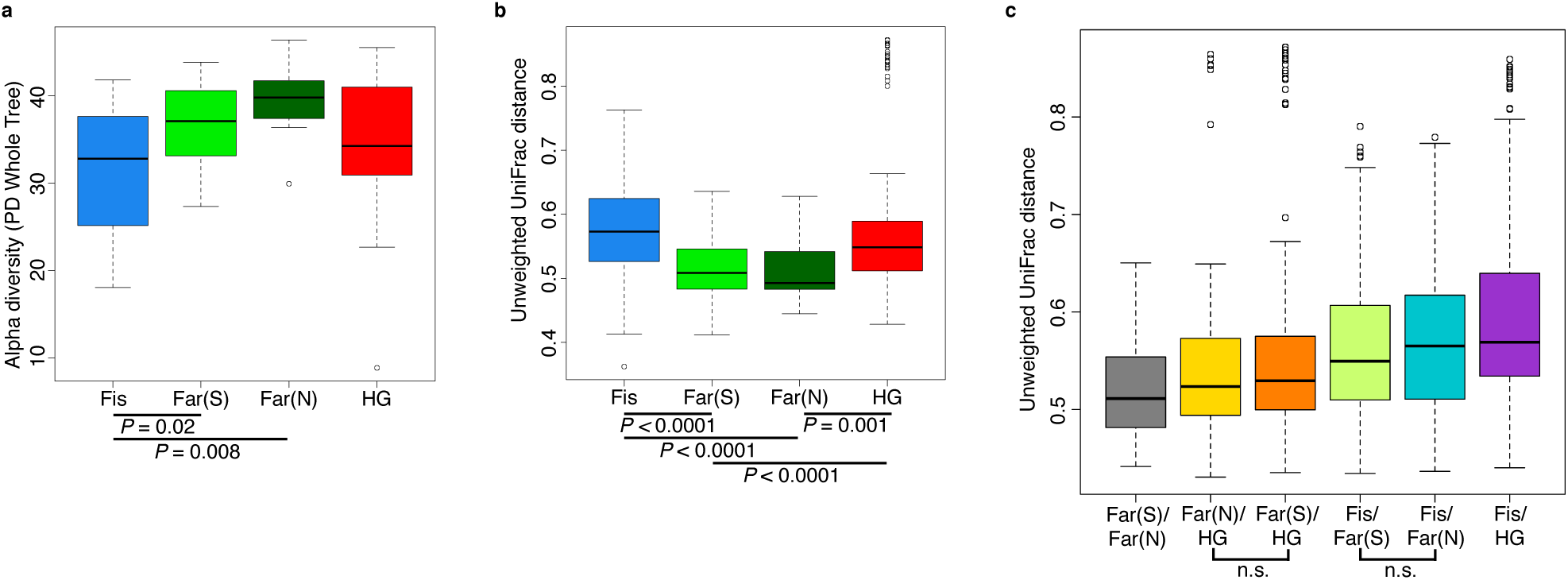
Comparison of the diversity of gut microbiomes of individuals across subsistence (a) Alpha diversity based on the phylogenetic metric, phylogenetic distance (PD) whole tree. (b) Beta diversity within each subsistence group based on unweighted UniFrac distances. (c) Beta diversity for pairs of subsistence groups based on unweighted UniFrac distances. For pairwise comparisons, all are significant (p < 0.05 unless specified (n.s.); Welch’s t-test). All p-values are based on Welch’s t-tests. Fis = Fishing population; Far(S) = Farmers from the South; Far(N) = Farmers from the North; HG = Hunter-gatherers.

Community differences between subsistence groups based on weighted and unweighted UniFrac distance metrics are the greatest between the fishing population and the other three subsistence groups (Figure 5c and Figure S17c). Beta diversity is the highest between the fishing population and hunter-gatherers, the two groups for which there is the largest number of significantly differentially abundant taxa (94% of significant taxa overall; Table S5). This finding might be expected, given that these populations differ not only in terms of diet, but also in their genetic ancestry and access to medicine. Both farmer populations are slightly more different from the fishing population than from the hunter-gatherers (Figure 5c and Figure S17c), and present a higher number of significantly differentially abundant taxa relative to the other populations (60% and 50% of the significant OTUs differ between the fishing population and the farmers from the South and the North, respectively, versus 44% and 38% that differ from the hunter-gatherers; Table S5). This could suggest that differences in access to medicine, or the occasional consumption of processed food in the fishing population, has considerable influence on their gut microbiomes, but this requires further investigation.

According to the taxonomy-based predicted metagenome for each subject’s gut microbiota, we found that only one pathway, bacterial invasion of epithelial cells, differed significantly across all subsistence types; represented at the highest relative abundance in the hunter-gatherers and lowest in the farmers (q = 0.03; ANOVA; Table S6 and Figure S18). This pathway includes proteins expressed by pathogenic bacteria that are important for entry into host cells. The importance of this difference is unclear, but could be indicative of an increased abundance of pathogens in the microbiomes of hunter-gatherers.

### Discussion

Here, we have investigated the relationship between intestinal parasitism and human gut microbiota, and found that presence of the protozoa *Entamoeba* is significantly correlated with gut microbiome composition and diversity across diet, geographic location, and genetic ancestry. Furthermore, we observed striking variation amongst different rural African populations despite a shared climate and ecosystem, indicating that there are multiple signatures of rural, unindustrialized microbiomes.

#### Influence of parasitism

The importance of gastrointestinal parasites in human disease is well established, both as infectious agents and in shaping immunity (30, 32, 54). This relationship is the basis of the hypothesis that the underlying cause of the high incidence of autoimmune diseases, unique to industrialized countries, is the absence of childhood exposure to infectious agents (36). Recent research supporting this hypothesis shows that mild and controlled infection by internal parasites can activate an immune response and reduce symptoms of a range of autoimmune diseases (55). Likewise, the relationship between gastrointestinal microbiota and host immune response has been well established (56-58). Despite evidence for direct host-parasite and host-microbiome interactions, and the fact that gut parasites and microbes share the same gut environment, studies are limited which assess the relationship between these organisms (31).

Here we show significant correlations between gut microbiota (composition and diversity) and the presence of the intestinal amoeba *Entamoeba (dispar, histolytica,* or both). Notably, individuals harboring these protozoa exhibit significantly higher alpha diversity in their bacterial gut communities, coupled with a significant reduction in inter-individual variation. This pattern could be a reflection of either direct or indirect interactions between *Entamoeba*, gut bacteria, and/or host immune factors. For example, *Entamoeba* could feed on certain species of bacteria, allowing others to proliferate or induce a host immune response that differentially affects the success of different microbes. Alternatively, it’s possible that a specific gut microbiota predisposes an individual to *Entamoeba* colonization. This relationship could also be the result of other correlating factors, not included in this study (e.g. exposure to anthelmintics and/or infection by other pathogens or parasites).

This pattern of lower alpha and higher beta diversity seen in *Entamoeba*-negative individuals has been repeatedly identified as a signature of gut microbiota in non-industrialized societies (15-18, 26). There are a number of hypotheses that have been proposed as explanations for this pervasive pattern including increased microbial dispersal (26), higher complexity of dietary carbohydrates (19, 22), and diminished or lack of exposure to antibiotics (59, 60).

An additional explanation for the inverse relationship between alpha and beta diversity is that in these populations, a more diverse gut microbiota is less sensitive to perturbations, or exhibits a limited number of potential stable states. It has been repeatedly demonstrated that biodiversity is often stabilizing, resulting in increased community resilience (61-63). However, there are exceptions to this common trend as exemplified in the people of Tunapuco, a traditional agricultural community from the Andean Highlands, who exhibit gut microbiota with both higher alpha and beta diversity compared to the Western community analyzed in the study (20). The explanation for this atypical pattern is not known, but it could be due to variables associated with the cooler climate of the region such as a distinct source community of microbes with more possible equilibrium states at higher levels of diversity and/or differences in parasite prevalence.

It is still unclear what mechanism is responsible for the observed differences in the gut microbiota of *Entamoeba* positive and negative individuals. We note that our study is only able to describe correlations between *Entamoeba* and the microbiome, and causality cannot be inferred. We expect further studies, perhaps using model organisms, to shed light on the causal factors underlying this relationship. However, these patterns are consistent across rural populations that vary in terms of geographic location, genetic ancestry, diet, and access to medicine, suggesting that the pervasiveness of intestinal parasites like *Entamoeba* in nonindustrialized societies might partially contribute to the explanation for the higher alpha diversity and lower beta diversity commonly observed in developing vs. industrialized populations. Alternatively, the relative differences in diversity between traditional vs. Western societies could be due to distinct and unrelated factors.

Interestingly, we found that the majority of specific taxa for which abundance significantly correlated with the presence of *Entamoeba* share the common feature that they have been highlighted for their potential role as signatures of inflammation-related diseases. For example, Clostridiales Ruminococcaceae, the second most important taxon in the RFC model, significantly more abundant in *Ent+* individuals, has been found to be underrepresented in individuals suffering from Crohn’s Disease and Ulcerative Colitis (27). Likewise, a decreased prevalence of *Prevotella copri* and Fusobacteria, as observed in *Ent*+ individuals, was recently shown to be negatively correlated with Rheumatoid arthritis (64) and incidence of colorectal cancer (65, 66), respectively. Although speculative, these relationships suggest a potential link between gut parasites, gut bacteria, and host inflammation. Additional studies are needed to elucidate the mechanisms driving these observed patterns, specifically how exposure to anthelmintics in developing countries might drive changes in gut microbiota that mirror patterns observed in industrialized societies.

#### Influence of subsistence and genetic ancestry

In addition to identifying the presence of *Entamoeba* as an important predictor of gut microbiome composition and structure across populations, we were also able to examine the relative influence of other factors. First, we compared the gut microbiome composition of individuals from the same subsistence mode and genetic ancestry, but coming from different villages. Within the hunter-gatherers, we saw clear differences in composition as well as in diversity between individuals living in Bandevouri versus those living in Makouré and Bidou, although this difference was only significant using the Shannon Index for alpha diversity (p < 0.05; Welch’s t-test). Based on the data we have available, we found that these groups do not differ in terms of diet or parasitism, suggesting a role for other unexplored very localized environmental factors (e.g., water source).

In terms of genetic ancestry, we found that, despite a genetic divergence as old as 60,000 years, the gut microbiome of the Pygmy hunter-gatherers is not strikingly different from that of the Bantu populations. The UniFrac distances and the number of significant taxa are indeed even lower between hunter-gatherers and farmers than between the farmers and the fishing population, two Bantu groups that share the same genetic ancestry. This suggests that in these populations, genetic background might play a smaller role in microbiome variation compared to the effect of diet and environment.

However, we found key differences distinguishing the microbiota of hunter-gatherers from those of the farming and fishing populations, likely reflecting the influence of their long-term diet. The hunter-gatherers were correctly assigned to their subsistence mode with higher accuracy (85%) relative to the other populations. Furthermore, some of our findings mirror patterns previously observed in comparisons of traditional vs. industrialized societies (18, 20, 26, 48) (see Table 1b), suggesting this ancestral subsistence mode might carry a specific microbial signature. Notably, we found a higher frequency of Proteobacteria in hunter-gatherers compared to the other Cameroonian populations, similar to the relationship between the Hadza and Italians (18) and that between traditional populations in Peru (hunter-gathers and farmers) and US individuals (20). Lachnospiraceae uncl, identified as the third important in the RFC model with the tendency to be lower in the Pygmy hunter-gatherers (5.7% versus 7.7-11.3% in other populations, q = 0.075), was also found to be in lower frequency in the Hadza compared to Italians (18). Finally, *Succinivibrio* and *Ruminobacter* species, enriched in the Hadza, were also identified as important taxa in the RFC model, and occur at higher frequencies in the Pygmy hunter gatherers (see Table 1b). Thus, all these taxa seem to be a specificity of hunter-gatherer populations, rather than reflecting a difference between industrialized European and rural African populations. *Succinivibrio* is considered to be an opportunistic pathogen, which could mean that hunter-gatherer populations have more opportunistic pathogens than other populations, as proposed by Schnorr et al (18). However, while *Treponema* was also found enriched in the Hadza, Matses, and Tunapuco populations (18, 20), we found it at a very low frequency in all the populations studied here (< 3.5%, Table 1b). Moreover, we found *Treponema* abundance to differ significantly based on *Entamoeba* infection status rather than subsistence. When looking at other opportunistic genera in the Enterobacteriaceae family, we found that *Shigella* and *Escherichia,* both previously found only in Italian children and not in children from Burkina Faso (15), occur at extremely low abundances in all four subsistence groups (< 0.1%, see Table 1b). As for *Klebsiella* and *Salmonella*, neither taxon differed significantly amongst our groups (Table 1b). Thus, there does not seem to be any clear trend for opportunistic pathogens in hunter-gatherers populations compared to others. However, our results highlight the importance of including parasite analysis in comparative studies of the gut microbiome of rural populations.

#### Influence of geography and industrialized lifestyle

Amongst the four populations included in this study, the fishing population is the most urbanized due to increased consumption of processed food and access to medicine. As such, the characteristics distinguishing the gut microbiomes of the fishing population from the farmers and hunter-gatherers that also differ between rural populations in developing countries and urban populations in industrialized countries (16, 18, 26) might correspond to signature patterns of a more industrialized lifestyle. In particular, within the phylum Bacteroidetes, we found a lower overall abundance of Bacteroidales uncl in the fishing population relative to the other three populations (Table 1), an order also depleted in Italians compared to Hadza (18). High abundance of *Prevotella* and *Bacteroides* have also been shown to represent signatures of the microbiomes for people in developing and industrialized countries, respectively (15, 16, 18, 20, 26). Higher abundances of *Prevotella* are often correlated with increased consumption of carbohydrates and simple sugars, whereas an elevated proportion of *Bacteroides* is associated with a diet richer in protein and fat. Although differences between the populations studied here were not statistically significant, the fishing population harbored the highest abundance of *Prevotella sp*. (30.8%), while the farmers from the North and the hunter-gatherers harbored the lowest (19.2% and 20.2%, respectively), congruent with decreased consumption of simple sugars in these populations. Notably, the abundance of *Prevotella sp.* is high relative to other genera in this order across all populations (Table 1b) and species of *Prevotella* were the most reduced in individuals infected with *Entamoeba*.

## Conclusion

This study suggests an important role for eukaryotic gut inhabitants and the potential for feedbacks between helminths, protozoa, microbes, and the host immune response, one that has been largely overlooked in studies of the microbiome. Prior analyses of the African gut microbiome have found an enrichment of *Treponema,* Bacteriodetes *a*nd *Prevotella* compared to European populations, an enrichment that has been proposed to be related to diet. However, our observations suggest that some of these trends could be related to the presence of *Entamoeba* (or other commensals and parasites). Furthermore, we found that many of the taxa for which abundance was significantly correlated with *Entamoeba* infection exhibit opposite patterns of abundance to those demonstrated to be correlated with a variety of autoimmune disorders. In addition, our results highlight the substantial variability in gut microbiome composition among closely related populations. Thus, using a single population as a representative of a lifestyle or geographical region may be overlooking important fine-scale patterns in microbiome diversity. Hence, comparative population studies of the human microbiome stand to benefit tremendously from considering variation within a geographic region and the role of parasitism and disease.

### Methods

**Sample collection.** We sampled 64 adult volunteers (26 females and 38 males) in seven rural villages (Bidou, Makouré, Bandevouri, Ndtoua, Afan Essokié, Akak and Ebodié) in Southern Cameroon, after obtaining their informed consent for this research project. The research permits, including all necessary ethical approvals, were obtained for this study by the “Institut de Recherche pour le Développement” (IRD) in agreement with the "Ministère de la Recherche Scientifique et de l’Innovation" (MINRESI) of Cameroon. For each participant, we collected information about his or her age, gender, anthropometric traits, health status, ethno-linguistic and quantitative nutritional questionnaires. We also collected saliva and fecal samples. The fecal sample was self-collected in the morning and stored in a plastic bag at most 3-4h before further handling. It was then split in two separate samples; one was used to perform the parasitological analysis at a local hospital (fresh or covered with formol) and the other was stored to run the sequencing analyses. This latter sample was handled following previous methods (67): the sample was first submerged with pure ethanol for about 24h at room temperature, then the ethanol was poured out of the container and the sample was wrapped in a sterile gauze and deposited on silica gel (18). The silica gel was then replaced by new gel when it changed colors from orange to yellow, i.e. when it could not absorb further humidity. The samples were then transported back to France and stored at -80°C until they were shipped to Minnesota, USA, on dry ice, and stored there at -80°C until further use. For five individuals, we were able to collect replicate fecal samples at two different time points: four individuals 7 days apart, and one individual 1 day apart.

### Characterization of intestinal parasitism

Intestinal parasitism of individuals was assessed by direct observation of fecal samples under the microscope. For each individual, a small amount of fecal matter was diluted in formol and homogenized to be liquid. A drop of liquid was then visualized under the microscope. If no known parasites were detected in this sample, another drop was closely inspected. If nothing was visible, the individual was characterized as negative. Parasite characterization was carried out by the same individual using the same method every time.

**DNA extraction and 16S rRNA amplification from the fecal sample.** Total DNA was extracted by bead beating from approximately 50 mg of each fecal sample using the MOBIO PowerFecal™ DNA Isolation Kit (MOBIO Laboratories, Carlsbad, CA, USA) according to the manufacturer's protocol. DNA isolated from fecal samples was quantified using a NanoDrop (ThermoScienctific), and the V5-V6 regions of the 16S rRNA gene were PCR amplified using Accura High Fidelity Polymerase, with the addition of barcodes for multiplexing. The forward and reverse primers were the V5F and V6R sets (68), chosen in part to allow dual coverage of the entire region. The barcoded amplicons were pooled and Illumina adapters were ligated to the reads. A single lane on an Illumina MiSeq instrument was used (250 cycles, 300 bp, pairedend) to generate 16S rRNA gene sequences yielding 175,784 Pass Filter (PF) reads per fecal sample (SD = 72,822) and ~12.65 million total PF reads (4.9Gb of data). Raw sequencing data (fastq files) are available through MG-RAST [MG-RAST ID pending].

**16S rRNA sequence analysis**. We obtained a total of 12.65 million high-quality reads, resulting in an average of 175,784 reads per sample (+/- 72,822). Raw Illumina sequences were demultiplexed and filtered using Cutadapt 1.7.1 (69) to remove adaptor sequences (Read 1:CTGTCTCTTATACACATCTCCGAGCCCACGAGAC, Read 2:CTCTCTCTTATACACATCTGCCGCTGCCGACGA), chimeras, sequences containing ambiguous bases, and low quality reads (Phred quality scores < 20). Read pairs were resynced using RISS-UTIL and matching paired-end sequences were merged using FLASH (70). Merged sequences over 250 bp in length (the maximum length of the V5-V6 region) were removed. The remaining merged sequences were analyzed using the open-source software package QIIME 1.7.0 (Quantitative Insights Into Microbial Ecology) (71). We performed both open- and closed-reference Operational Taxonomic Unit (OTU) picking at 97% identity against the May 2013 Greengenes database (72) such that OTUs were assigned taxonomy based on 97% similarity to the reference sequence. Non-bacterial 16S rRNA sequences removed and those that did not align were clustered to each other prior to taxonomic assignment. The average percent of mapped reads per individual was 83% (SD = 7.5%) and did not vary significantly between populations (Welch’s t-test, p > 0.2). All summaries of the taxonomic distributions ranging from phylum to species were generated from the non-rarefied OTU table generated from this analysis.

### Diversity analyses

To characterize diversity across individuals, rarefaction plots were generated for each sample using the phylogenetic distance metric for diversity (73). Samples were rarefied to 50,000 reads, the maximum depth permitted to retain all samples in the dataset. All diversity analyses were conducted on rarefied OTU tables containing 50,000 sequences per sample. Measurements are based on the mean values calculated from 100 iterations using a rarefaction of 10,000 sequences per sample (20% of the total 50,000). Alpha-diversity was calculated for each sample based on phylogenetic diversity, Shannon’s index (74) and the Simpson index (75). Beta-diversity was assessed based on both unweighted and weighted UniFrac distance metrics (43) using the Greengenes phylogenetic tree (72). Principal Coordinate Analysis (PCoA) was carried out on the distance matrices. P-values were calculated using the Welch’s or Wilcoxon ttests, depending on normality of the distribution. To determine if the UniFrac distances were on average significantly different for groups of samples, we conducted Principal Coordinates Analysis (PCoA) to reduce raw gastrointestinal microbial community data into axes of variation. We assessed the significance of each covariate by performing a permutational multivariate analysis of variance (PERMANOVA) (76), a non-parametric test, on both weighted and unweighted UniFrac distance matrices using the “adonis” function from the *vegan* package in R (77). This test compares the intragroup and intergroup distances using a permutation scheme to calculates a p-value. For all PERMANOVA tests we used 10,000 randomizations.

### Multivariate analysis of composition data

Intergroup differences in microbiome composition for subsistence, location, population, BMI, sex, ancestry, age, dietary factors, and parasitism were assessed by PERMANOVA (76) using nonrarefied OTU abundance data and implemented using the “adonis” function of the *vegan* package in R (77). To identify taxa significantly associated with each covariate of interest, we normalized the distribution of each OTU and used an ANOVA, FDR corrected for the number of OTUs in our dataset. For the ANOVA, OTUs with identical taxonomic identifiers were combined. In parallel, we also restricted the merging only to OTUs names defined at the family, genus or species level. Both results are reported in Supplementary Material (“merged OTUs” versus “partially merged OTUs”). For analyses of both merged and partially merged OTUs, the resulting taxa were filtered to include only those that occurred at least 0.1% in at least 4 individuals.

### Random forest classifier model

A random forest classifier with 2000 decision trees was trained on the taxa abundance table consisting of 93 OTUs with 5-fold cross-validation using scikit-learn (78). Mean accuracy (the ration of the number of correct predictions relative to the total number of predictions) over the 5 folds was 0.79 (standard deviation 0.09) with p < 0.001 (estimated using 1000 permutation tests with 5-fold cross-validation). The most discriminating taxa were identified by random forest importance values (in scikit-learn random forest importance values are calculated as mean decrease in node impurity). We report the top ten median importance values and 95% confidence intervals from 1000 random forests.

### Metagenomic predictions

We used PICRUSt v1.0.0 (Phylogenetic Investigation of Communities by Reconstruction of Unobserved States) to generate taxonomy-based predicted metagenomes for each sample (50). Counts from the rarefied OTU Table (50,000 OTUs per sample) were normalized by the predicted 16S rRNA gene abundances and functional predictions of Kyoto Encyclopedia of Genes and Genomes (KEGG) (49) pathways were determined using pre-computed files for the May 2013 Greengenes database (72). Relative abundances of the functional predictions were calculated. We also compared the predicted metagenomes of individuals to determine which functions were enriched or depleted across covariates (subsistence, population, location, ancestry, BMI, age, sex, dietary components, and parasitism phenotypes) for abundant (≥ 0.1% in at least 4 individuals) and rare (< 0.1% in at least 4 individuals) pathways. An ANOVA was used to determine which predicted pathways were significant (q < 0.05) for each covariate.

## Acknowledgements

We thank people on the field that volunteered for this project as well as those that helped for the collection of data. We also thank Karen Tang, Andres Gomez, Michael Burns, and Peter Zee for helpful discussions. This work was carried out using computing resources at the Minnesota Supercomputing Institute.

## Related Manuscripts

The authors have no related work under consideration for publication at this time.

